# Universal Ontogenetic Growth without Fitted Parameters: Implications for Life History Invariants & Population Growth

**DOI:** 10.1101/2021.10.10.463814

**Authors:** Andrés Escala

## Abstract

Since the work of Von Bertalanffy (1957), several models have been proposed that relate the ontogenetic scaling of energy assimilation and metabolism to growth, which are able to describe ontogenetic growth trajectories for living organisms and collapse them onto a single universal curve (West et al. 2001; Barnavar et al. 2002). Nevertheless, all these ontogenetic growth models critically depend on fitting parameters and on the allometric scaling of the metabolic rate. Using a new metabolic rate relation (Escala 2019) applied to a Bertalanffy-type ontogenetic growth equation, we find that ontogenetic growth can also be described by a universal growth curve for all studied species, but without the aid of any fitting parameters (i.e., no fitting procedure is performed on individual growth curves). We find that the inverse of the heart frequency f_H_, rescaled by the ratio of the specific energies for biomass creation and metabolism, defines the characteristic timescale for ontogenetic growth. Moreover, our model also predicts a generation time and lifespan that explain the origin of several ‘Life History Invariants’ (Charnov 1993) and predict that the Malthusian parameter should be inversely proportional to both the generation time and lifespan, in agreement with the data in the literature (Duncan et al. 1997; Dillingham et. al 2016; Hatton et al 2019). In our formalism, several critical timescales and rates (lifespan, generation time and intrinsic population growth rate) are all proportional to the heart frequency f_H_, and thus, their allometric scaling relations come directly from the allometry of the heart frequency, which is typically f_H_ ∝ M^−0.25^ under basal conditions.

## 1. Introduction

Metabolism and growth are two fundamental aspects of living organisms, and thus, it is somewhat natural to try to understand the connections between these two processes. Von Bertalanffy (1957) studied ontogenetic growth curves in order to establish connections between metabolism and growth, showing that individual growth curves in living organisms can be reproduced from models of metabolic energy allocation. Although several subsequent growth models (Reiss 1989; West et al. 2001; Ricklefs 2003; Hou et al. 2008) differ significantly on the details of their derivations, they share the same mathematical form of Von Bertalanffy (1957), which basically links patterns of assimilation and growth to the (allometric) scaling of the metabolic rate. In particular, West et al. (2001) proposed a general quantitative model based on the allocation of metabolic energy and showed that individual growth curves can be collapsed onto a single universal curve that describes the growth in all the studied species.

Nevertheless, Banavar et al (2002) later illustrated that the universal growth curve arises from general considerations of energy conservation that are independent of the specific allometric model used by West et al. (2001). In particular, Banavar et al (2002) showed that the data do not distinguish between specific exponents in the scaling relationship between metabolic rate and mass, with exponents of 2/3 and 3/4 in the metabolic relation fitting equally well. This scaling collapse onto a universal curve, which occurs when some dimensionless quantities are properly defined, is also equivalent to asserting that a single self-similar solution is able to successfully fit all the ontogenetic growth curves. Since the universal curve arises from general considerations, it is desirable to find an independent test of the assumptions behind the models that defines the key dimensionless quantities that can discriminate between the different models for ontogenetic growth.

In general, all ontogenetic growth models critically depend on the metabolic rate, specifically, on the allometric scaling, the exact slope of which is still matter of debate (White et al. 2007). Recently, the empirical metabolic rate relation was corrected in order to fulfill dimensional homogeneity (Escala 2019), a minimal requirement for any meaningful law of nature (Bridgman 1922), and a new metabolic rate (B) formula was proposed: B = *ϵ*(T) *η*_O_2__ f_H_ M, where M is the body mass, f_H_ is a (characteristic) heart frequency, *η*_O_2__ is a specific O_2_ absorption factor for different exercising conditions (basal, maximal, etc.) and *ϵ*(T) = *ϵ*_0_ e^−E_a_/kT^ is a temperature correction inspired by the Arrhenius formula (Gillooly et al. 2001), in which E_a_ is the activation energy and k is the Boltzmann constant. Compared to Kleiber’s original formulation (Kleiber 1932), B = B_0_(M/M_0_)^0.75^, and Rubner’s surface rule of proportionality to 2/3, this new metabolic rate relation has the heart frequency f_H_ as the controlling variable (a marker of metabolic rate); its advantage is that it is a unique metabolic rate equation for different classes of animals and different exercising conditions that is valid for both basal and maximal metabolic rates, in agreement with empirical data in the literature (Escala 2019). In addition, Escala (2022) showed that this new metabolic rate relation can be directly linked to the total energy consumed in a lifespan, so it is able to explain the origin of variations in the ‘rate of living’ theory (Speakman 2005; Ramsey et al. 2000).

In this paper, we explore the implications of this new metabolic relation for ontogenetic growth models, with a focus on independently testing key quantities in our formulation to enable discrimination between previous models. The paper is organized as follows: We start in §2 by applying the results of the new metabolic rate relation (Escala 2019) to an ontogenetic growth equation that shares the same mathematical form as previous models, showing that individual growth curves can also be collapsed onto a single universal curve, but in this case without the aid of any fitting parameter. Section 3 continues by computing the predicted generation time and explaining the origin of several ‘Life History Invariants’, with satisfactory results. In §4, we study the predicted implications for population growth, showing that they agree with the collected data. Finally, in §5, we discuss the results and final implications of this work.

## 2. Ontogenetic Growth Model

Since different assumptions about energy allocation lead to the same general equation (von Bertalanffy 1957; Reiss 1989; West et al. 2001; Ricklefs 2003; Hou et al. 2008), we will follow the notation in the mass-energy conservation model described in Moses et al. (2008), which revisited the ontogenetic growth model of West et al. (2001), because we will compare our results to theirs, using their parameter estimations. The conservation of energy for the allocation of metabolic energy during growth between the maintenance of existing tissue and the production of new biomass can be expressed as 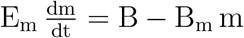 (Moses et al. 2008), where B is the metabolic rate (in J/s or W), E_m_ (in J/g) is the energy required to create a unit of biomass and B_m_ (in W/g) is the metabolic rate required to maintain an existing unit of biomass.

For the corrected metabolic rate relation, we will restrict ourselves to basal (resting) conditions and neglect temperature variations because ontogenetic growth happens over long periods of time during which such variations might tend to cancel each other out (in wild conditions). Under such conditions, we have the constant factor *ϵ*(T) *η*_O_2__ ≈ 10^−4.313^ mlO_2_g^−1^ ≈ 10^−3^ J/g ≡ E_2019_ (converting 1 ltr O_2_=20.1 kJ; Schmidt-Nielsen 1984), where E_2019_ is a constant that comes from the best fitted value for the corrected metabolic relation (Eq. 8 of Escala 2019). Therefore, the metabolic rate formula is simply given by B = E_2019_f_H_ m (Escala 2019), and assuming that the heart frequency scales with body mass m as f_H_ = f_0_m^−*α*^, (mass-)energy conservation can be expressed as

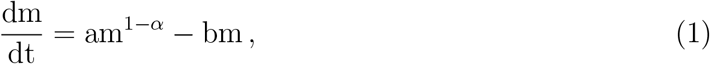

where a = E_2019_ f_0_/E_m_ and b = B_m_/E_m_. The general solution of Eq. 1 is a classical sigmoidal curve, the general form of which was given by von Bertalanffy (1957; Eq. 6). Noting also that for an initial (birth) mass *m*_0_ and final (asymptotic) mass M, the condition dm/dt = 0 (at m = M) in Eq. 1 is equivalent to a/b = M^*α*^, the solution to Eq. 1 can be written as:

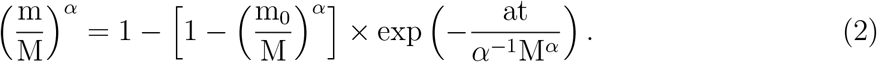

This solution is equivalent to the one found by Banavar et al. (2002), and for the special case of *α* = 1/4, it has the solution given in West et al (2001) (their Eq. 5). The solution given by Eq. 2 can be rewritten as r = 1 – e^−*τ*^, which is the same universal growth curve found in West et al (2001) (and Banavar et al 2002), but in our case, it uses the following variable change:

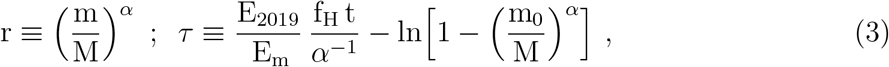

where we also replace a = E_2019_ f_0_/E_m_ in Eq. 2 (in the definition of *τ*), and thus, f_H_ = f_0_M^−*α*^ is (henceforth) the heart frequency when the animal reaches the final (asymptotic) mass M. For the particular case of *α* = 1/3, r also corresponds to the fractional size (r = (m/M)^1/3^ = *l*/L), leading to the classical Bertalanffy growth equation, *l*(*τ*) = L(1 – e^−*τ*^), which has been successfully applied in the fishery industry (Beverton & Holt, 1959; Charnov 2008). In this case, we can explicitly express (in terms of physical quantities) the Bertalanffy growth coefficient as 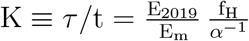 (from Eq. 3 for m_0_ = 0).

The advantage of this formulation compared to previous ones is that now the dimensionless time *τ* is expressed exclusively in terms of quantities that are well defined, with either clear physical (specific energies E_2019_ and E_m_) or biological (heart frequency f_H_, body mass M, etc.) meanings, and therefore, the dimensionless variables defined in Eq. 3 are now written in a physically transparent form, without the aid of any fitting parameters. This contrasts, for example, with the parameter a (present in Eqs. 1 & 2 but not in the universal growth curve defined by Eq. 3) that is critical in the procedure of fitting the growth curves in Bertalanffy (1957), West et al (2001), Banavar et al (2002), etc. and in the definition of their dimensionless variables, which has an obscure meaning considering its fractal dimensionality of [mass^*α*^]/[time] and is thus not associable to any physical quantity. This fitting parameter a is not present in the final growth curve solution found in our formulation, r = 1 – e^−*τ*^, for the variables defined in Eq. 3. It is notable that in this formulation for *τ*, the inverse of the heart frequency 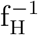 defines the characteristic timescale for ontogenetic growth, 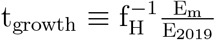, which is rescaled by the ratio of the specific energies of biomass creation (E_m_) and (basal) metabolism (E_2019_).

Banavar et al. (2002) showed that the ontogenetic growth curves do not distinguish between exponents of 2/3 and 3/4 (= 1 – *α*) in the mass scaling of the metabolic rate relation, as both are equally good fits to the current data and are consistent with a universal curve r = 1 – e^−*τ*^ for ontogenetic growth. Therefore, only for consistency, we will assume *α* = 1/4, since we will compare our formulation against data using the fitting parameters from West et al (2001), which assumes this *α* value.

The universal growth curve r = 1 – e^−*τ*^, for the *τ* defined in Eq. 3 (with *α* = 1/4), is mathematically identical to the solution found in West et al (2001) (their Eq. 5) when

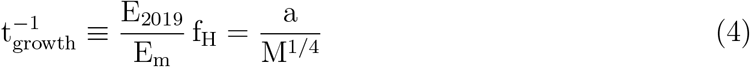

is fulfilled, and therefore, it should equally well fit the individual growth curves for the species studied in West et al (2001) if their measured heart frequencies f_H_ and estimated specific energies (E_2019_, E_m_) fulfill the condition given by Eq. 4 for the parameter a and final mass M found in the fitting procedure of the growth curves in West et al (2001). In order to test whether the condition in Eq. 4 is fulfilled in nature, we collected estimations for the energy required to create a unit of biomass E_m_ (Moses et al. 2008, taking averages when multiple estimations are available) and measurements of heart frequencies f_H_ for the species studied in West et al (2001) in order to supplement them with the fitting parameters a, birth mass m_0_ and final mass M that best fit the growth curves studied in West et al (2001). A summary of all the quantities (with the individual references for f_H_) are listed in Table 1.

**Table 1.**

| Species | $a$ [$\text{g}^{1/4}/\text{day}$] | $m_0$ [kg] | $M$ [kg] | $E_m$ [J/g] | $f_H$ [# / min] |
| --- | --- | --- | --- | --- | --- |
|  |  | West et al 2001 |  | Moses et al. 2008 | (Reference) |
| Cow | 0.28 | 33.3 | 442 | 6950 | 63 (Kovacs et al 2005) |
| Pig | 0.31 | 0.90 | 320 | 6150 | 70 (Harris 2009) |
| Rabbit | 0.36 | 0.12 | 1.35 | 6100 | 180 (Detweiler et al. 2004) |
| Guinea pig | 0.21 | 0.005 | 0.840 | 4900 | 277 (Weibel et al. 2004) |
| Rat | 0.23 | 0.008 | 0.280 | 4350 | 393 (Weibel et al. 2004) |
| Shrew | 0.83 | 0.0003 | 0.0042 | 1800 | 835 (Jurgens et al. 1996) |
| Heron | 1.56 | 0.003 | 2.7 | 1400 | 211 (Machida et al. 2001) |
| Robin | 1.9 | 0.001 | 0.022 | 2000 | 570 (Welty et al. 1988) |
| Cod | 0.017 | 0.0001 | 25 | 13000 | 32 (Wardle et al. 1973) |
| Salmon | 0.026 | 0.00001 | 2.4 | 7300 | 30 (Clark et al. 2011) |
| Guppy | 0.10 | 0.000008 | 0.00015 | 1900 | 120 (Unpublished data) |

Figure 1 displays the relation given by Eq. 4 between the variables a, M, E_m_ & f_H_ as a function of the final body mass M. In addition to the 7 orders of magnitude in body mass M variations, the relation has a slope consistent with zero (∝ M^−0.05^, and there is only 0.22 dex in scatter, where 1 dex on a log scale refers to an order of magnitude, comparable to the scatter in the metabolic rate relation found in Escala 2019), confirming that the condition in Eq. 4 is fulfilled. This implies that for the specific dimensionless variables defined in Eq. 3, the universal curve (r = 1 – e^−*τ*^) will describe the (different) individual growth curves as well as the models in West et al (2001) (and Banavar et al. 2002). This is found without performing any fitting procedure on individual growth curves; instead, it is derived from the mathematical equivalence between our universal growth curve (r = 1 – e^−*τ*^, for the *τ* defined in Eq. 3) and West et al (2001)’s solution when Eq. 4 is fulfilled. In other words, for the same universal solution, our formalism replaces the fitting parameters in West et al (2001), such as a, which involves obscure units [g^1/4^/day] and is thus not easily associable to a known physical quantity, by well-defined energies (E_2019_ & E_m_) and frequency (f_H_), which are also determined independently rather than by fitting animal growth curves (as in West et al 2001, Banavar et al. 2002, etc.).

**Fig. 1.**
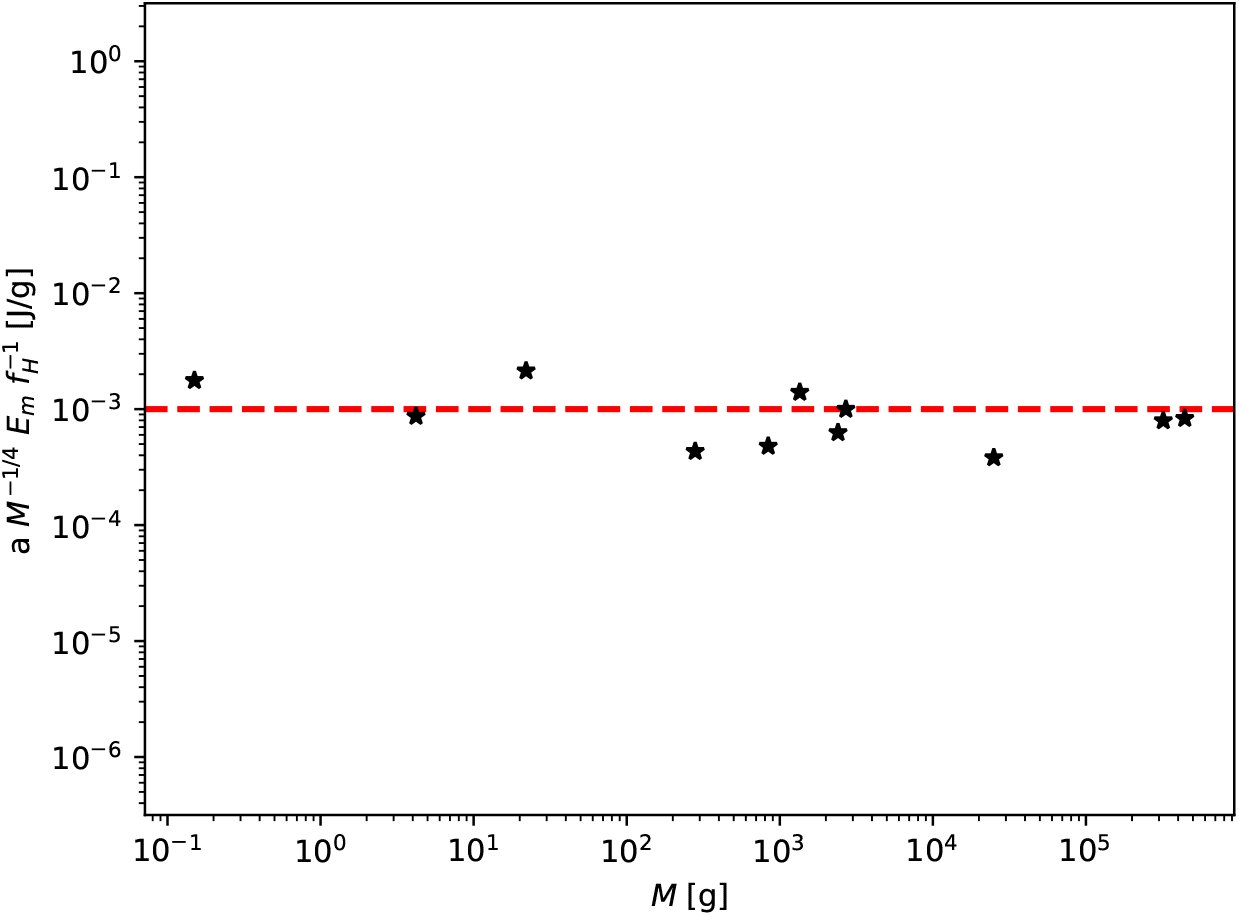
The relation between the variables a, M, E_m_ & f_H_, as a function of the final body mass M, showing that it does not vary systematically with M, as predicted by Eq. 4. The different species marked with black stars are scattered around the predicted value of E_2019_, which is denoted by the red dashed line. The best-fitted slope is 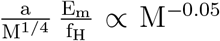, which is consistent with a zero slope against body mass M. The parameters a and M, taken from West et al (2001), are determined to best fit the individual growth curves of the different species. In contrast, the parameters E_2019_, E_m_ & f_H_ that define t_growth_ in our formulation are determined independently (see the references in Table 1) from any procedure of fitting to individual growth curves, such as the one in West et al (2001).

Moreover, the different species shown in Figure 1 are scattered around the predicted value of E_2019_ (= 10^−3^ Jg^−1^; Escala 2019) denoted by the red dashed line, and therefore, Fig 1 can be considered a third independent estimation of E_2019_ (in addition to those of Escala 2019, 2022), supporting the predictability of the formalism presented and showing a characteristic feature of the physical sciences: fewer (and more universal) constants that have consistent measurements in multiple contexts. Finally, Figure 1 strongly supports the idea that the inverse of the heart frequency 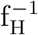, rescaled by the ratio of the competing specific energies (biomass creation E_m_ versus metabolism E_2019_), defines the characteristic timescale for ontogenetic growth 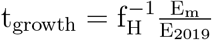.

## 3. Generation Time and the Origin of Some Life History Invariants

The success of the formulation given by Eq. 3, supported by Fig 1, motivates us to study its implications, and for that reason, it is interesting to look at the predicted generation time t_gen_, which is the time period required for a young organism to grow to its final size and thus mature to reproductive age. The generation time t_gen_ can be straightforwardly determined from the *τ* defined in Eq. 3, making it possible to arrive at the following relation:

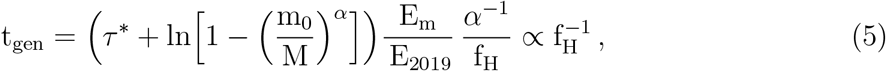

where *τ*\* is the approximate dimensionless time value for reaching adult mass (for all species) in the universal growth curve. Eq. 5 gives a generation time that is explicitly dependent on the heart frequency f_H_, where the allometric scaling of the heart frequency under basal conditions (f_H_ ∝ M^−0.25^; Brody 1945, Calder 1968) gives the well known mass scaling for the generation time (t_gen_ ∝ M^0.25^; Bonner 1965). Nevertheless, since most biological rates and times scale as M^−1/4^ and M^1/4^ (Savage et al. 2004, Burger et al. 2021), an interesting possible test is to study allometric variations of t_gen_ in large outliers of 1/4 scaling such as spiders (Anderson 1970, 1974) (White et al. 2007) to test whether they vary as f_H_, as predicted by Eq. 5. Another interesting possibility is to directly test the predicted inverse correlation between t_gen_ and f_H_.

One of the advantages of having a generation time that is explicitly dependent on the heart frequency f_H_is that it can be directly linked (Escala 2019, 2022) to the total lifespan t_life_ using the empirical relation of the total number of heartbeats (N_b_) in a lifetime t_life_ = N_b_/f_H_ found to be valid in mammals (Levine 1997; Cook et. al 2006). Escala (2022) generalized this relation for all types of living organisms using the proportionality between heart and respiration frequencies, f_H_ = kf_resp_ (Schmidt-Nielsen 1984); then, the empirical relation with lifetime can be rewritten so that it is also valid for a total number N_r_ (= N_b_/k) of ‘respiration cycles’: t_life_ = N_b_/f_H_ = N_b_/kf_resp_ = N_r_/f_resp_. Escala (2022) also used this relation to predict the total lifespan energy consumed in living organisms, with satisfactory results if an approximately constant total number N_r_ ~ 10^8^ of respiration cycles per lifetime is required in all living organisms, which further supports the generalization. Combining this relation for a fixed number of respiration cycles per lifetime with Eq. 5, we obtain:

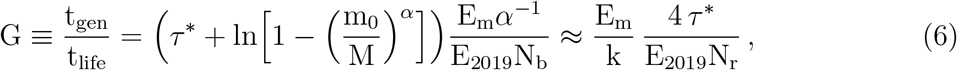

where we also approximate m_0_ ≪ M (in agreement with the species listed in Table 1), neglecting minor changes due to the weakly varying logarithm. Eq. 6 directly relates the generation time and lifespan, two quantities that are known to correlate (de Magalhaes et al 2007) with their respective energies (per unit mass) to create biomass (E_m_) and sustain lifespan (N_b_E_2019_), by only assuming (mass-)energy conservation (Eq. 1) and the invariant N_r_ (= 1.62 10^8^ respiration cycles per lifetime; Escala 2022) as a critical link between the two timescales.

Eq. 6 can be directly compared to the data compiled by Charnov & Berrigan (1990), which summarized published data for the ratio t_adult_/t_gen_ in different animal groups, where t_adult_ = t_life_ – t_gen_ is the adult lifespan, noting that this definition is related to the G ratio as t_gen_/t_life_ = (1 + t_adult_/t_gen_)^−1^. Additionally, we note that in Charnov & Berrigan (1990), lifespans are estimated from the inverse of (instantaneous) mortality rates, being effectively field lifespans and not maximum ones; thus, for comparison with Eq. 6, a correcting factor of 2.5 must be included (McCoy & Gillooly 2008), namely, 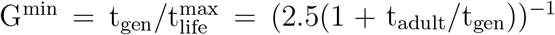.

Table 2 summarizes the estimations of G^min^ using the data compiled in Charnov & Berrigan (1990) and compares them with the predicted values from Eq. 6, using estimates of E_m_ for taxonomic groups (average of juvenile estimates; Moses et al. 2008), k values for such groups (Schmidt-Nielsen 1984; Escala 2022) and a dimensionless time *τ*\* ~ 5, the approximate value to reach adult mass in the universal growth curve (i.e., the flat part of the curve in Fig 2 of West et al 2001). Overall, we find that the values predicted from Eq. 6 are consistent with the empirically determined ones, as can be seen in Table 2. This is in addition to the consideration that each taxonomic group of birds, mammals and fishes includes species with up to an order of magnitude difference in E_m_ values (i.e., cod vs guppy fish in Table 1) and not necessarily the same species studied in Charnov & Berrigan (1990). Therefore, it is even more relevant that the relative G^min^ trends (between birds, mammals and fish) observed in Charnov & Berrigan (1990) are successfully predicted by the ratio 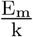.

**Fig. 2.**
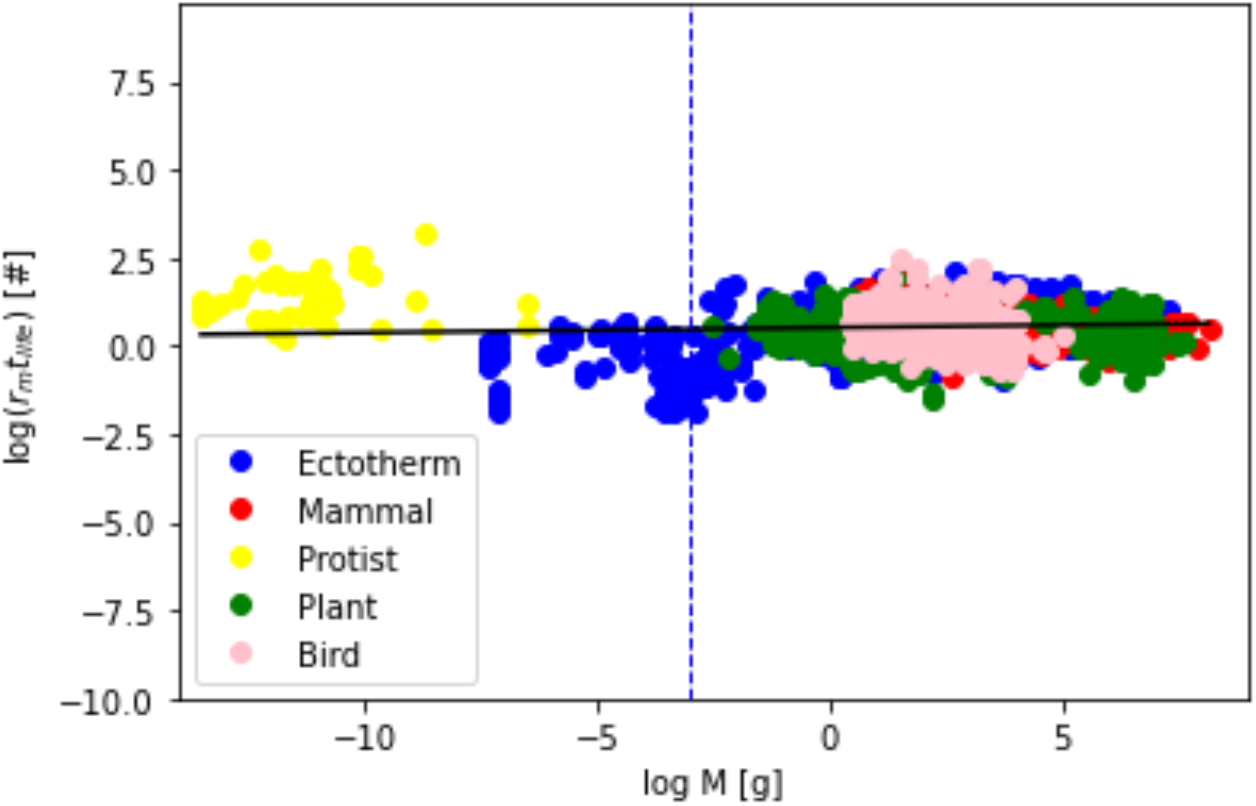
Lifetime Growth: Intrinsic (maximum) population growth rates (r_m_) multiplied by the corresponding lifespans (t_life_) for all types of living organisms, ranging from protists to mammals and birds, compiled by Hatton et al (2019). The solid black line corresponds to the best fit to the data, with no residual slope with body mass, r_m_t_life_ = 3.35 M^0.01^, as expected from Eq. 8, with a scatter of 0.5 dex. The dashed blue line denotes the mass scale of 10^−3^ grams, which corresponds to the scale associated with the smallest organisms with hearts, to which the present analysis is restricted.

**Table 2.**

| Animal Group | $E_m$ [J/g] | k | Predicted from Eq. 6 | $G^{\min} = \frac{t_{\text{gen}}}{t_{\text{life}}^{\text{max}}}$ |
| --- | --- | --- | --- | --- |
| Birds | 4050 | 9 | 0.06 | 0.11 |
| Mammals | 5800 | 4.5 | 0.16 | 0.17 |
| Fish | 7400 | 3 | 0.3 | 0.28 |

Eq. 6 shows that the invariant relation between lifespan and age at maturity that is valid within taxonomic groups, as observed by Charnov & Berrigan (1990), might come directly from the existence of another invariant: the approximately constant total number N_r_ ~ 10^8^ of respiration cycles per lifetime. This ratio t_adult_/t_gen_ is indeed one of the studied dimensionless life-history invariants (Calder 1984; Charnov 1993), which has sometimes been criticized as spurious by being a form of ‘regressing X on X’, where X is a random number (Nee et al. 2005). In this work, the generation and lifespan timescales are quantities derived independently, giving a quantitative prediction of an approximately invariant ratio within taxonomic groups, including interspecific variations, that comes from the basic energetics of respiration and the creation of new biomass and therefore has a clear physical interpretation and is thus far from spurious.

Traditionally, this relation, expressed in terms of the adult lifespan and age at maturity (which typically marks the end of an animal’s growth), as in Charnov & Berrigan (1990), has been qualitatively explained in terms of life-history evolution theory. Many versions of life-history theories predict that the age of maturity should be positively correlated with the lifespan (Charnov 1993), as these patterns are a reflection of natural selection (Charnov & Berrigan 1991). In this paper, after rewriting the properties of living organisms in a physically transparent form, we predict the value of this life-history invariant in terms of the relevant energetics (E_m_, E_2019_, k, etc.) and find that the constancy mainly comes from the invariant number N_r_ ~ 10^8^ of respiration cycles per lifetime, a generalization of the well-known relation of a constant number of heartbeats in a mammal’s lifetime (Levine 1997; Cook et. al 2006). Nevertheless, we do not study the origin of how the key parameters (E_m_, k, etc.)vary across species, animal groups and generations, which should be evolutionary in origin and thus set by natural selection (Charnov 1993; Gardner et al 2005).

It can be straightforwardly obtained that some of the other life-history invariants (Charnov 1993) come directly from the invariant G = t_adult_/t_gen_ and the universal ontogenetic growth curve. For example, the invariant ratio between the mortality rate (M) and the Bertalanffy growth coefficient (K) (Beverton & Holt 1963, Cushing 1968, Pauly 1980) comes from its relation with τ in our formulation, Kt = *τ*, which at the generation time corresponds to Kt_gen_ = *τ*\* ~ 5 according to the universal growth curve (West et al. 2001; Barnavar et al. 2002). Noting also that the mortality rate M is approximately the inverse of the lifespan 1/tiife (in the next section, we will see a more rigorous definition for M), it is straightforward to obtain that M/K = t_gen_/(t_life_*τ*\*) = G/5. Another invariant is the ratio between the length at maturity *l*(t_gen_) and the maximum asymptotic length L (Charnov & Berrigan 1991), which also comes from Kt_gen_ = *τ*\* ~ 5 by simply recalling the Bertalanffy growth equation (derived in §2): l (t_gen_)/L = 1 – e^−*τ*^*. Finally, the invariant fraction of body mass to reproduction per unit time per life span (Charnov, Turner & Winemiller 2001) comes from the definition of the dimensionless mass r (Eq. 3) and the invariant G (Eq. 6), which are related as = (1 – e^−*τ*^*)^1/*α*^ G^−1^, noting again that *τ*\* ~ 5.

## 4. Implications for Population Growth: The Malthusian Parameter and the ‘Equal Fitness Paradigm’

Malthus (1798) studied the simplest model of population growth, which can be derived by assuming that all individuals are identical and reproduce continuously; therefore, the number of individuals N will change with the birth rate B and death rate D as follows:

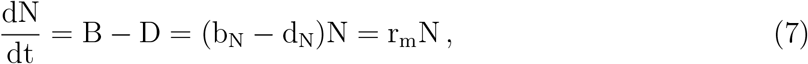

where b_N_ and d_N_ are the per capita birth and death rates, respectively, and r_m_ is the Malthusian parameter or intrinsic (maximum) population growth rate. The solution of Eq. 7 has an exponential form given by N(t) = N_0_ e^r_m_t^, and therefore, the Malthusian parameter r_m_ = b_N_ – d_N_ has units of inverse time and can be rewritten in the following form: 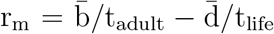. The constants 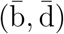 are now dimensionless since we have identified t_life_ as the characteristic timescale for death rates and t_adult_ = t_life_ – t_gen_ as the characteristic timescale for the birth rates. The reason for the latter is that t_adult_ is the reproductive adult lifespan, which is determined by subtracting from the lifespan the time period required to (grow and) mature to reproductive age, t_gen_, which does not fulfill the key assumption in Eq. 7 that the individuals are able to reproduce continuously.

Using Eq. 6, which relates t_gen_ and t_life_, the Malthusian parameter is given by:

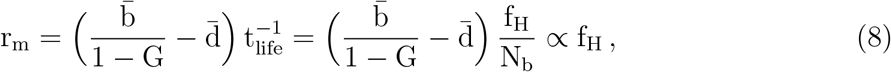

which gives a Malthusian parameter r_m_ that is again explicitly dependent on the heart frequency f_H_. The allometric scaling of heart frequency under basal conditions (f_H_ ∝ M^−0.25^, Brody 1945, Calder 1968) gives the well-known Fenchel (1974) allometry, r_m_ ∝ M^−0.25^, but again, as most biological rates and times scale as M^−1/4^ and M^1/4^ (Savage et al. 2004, Burger et al. 2021), a more interesting test will be to study allometric variations of r_m_ in outliers of ‘1/4 scaling’ in terms of the heart frequency f_H_ to determine whether they vary as f_H_, as predicted by Eq. 8.

Another prediction of Eq. 8 is an inverse correlation between r_m_ andt_life_, which was recently tested by Hatton et al (2019). Fig 2 displays the data of Hatton et al (2019) for all types of living organisms, ranging from protists to mammals and birds. Since the present analysis is restricted to organisms with a heart, we denote with the dashed (blue) line the mass scale of 10^−3^ grams, which corresponds to the scale associated with the smallest organisms with hearts, such as fruit and fairy flies, and coincides in Fig 2 with a transition to a decrease in the scatter (for larger masses); in addition, all individual animal groups (colors in Fig 2) satisfy this relation for masses larger than 10^−3^ grams. The solid black line corresponds to the best fit to the data, consistent with a clear inverse correlation between r_m_ and t_life_, with no residual slope with body mass (r_m_t_life_ = 3.35 M^0.01^, scatter of 0.5 dex or half an order of magnitude).

The relatively low variation (on average, one order of magnitude spread, since the scatter is 0.5 dex) for a database that includes sources of scatter such as mixing lifespans in captivity and in the wild (Hatton et al 2019; see de Magalhaes & Costa 2009 for more on this issue) and in particular a negligible slope with body mass as shown in Fig 2 has been claimed (Hatton et al 2019) to support the so-called ‘Equal Fitness Paradigm’ (Brown et al 2018; Burger et al. 2020), which states that “most organisms are more or less equally fit, as evidenced by the persistence of millions of plant, animal and microbe species of widely varying size, form and function in the Earth’s diverse environments” (Brown et al 2018). In terms of the proposed formalism, Eq. 8 and the low variation of G across animal groups (the factors 2-3 seen in Table 2), also implies a low variation for the parameters 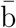 and 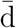 across species. These relatively constant values (implied by Fig 2) for the dimensionless parameters 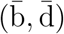 associated with the (net) per capita birth and death per characteristic time, in addition to the spectacular diversity of life histories in terms of growth, reproduction and survivorship over the life cycle, which implies very diverse life history strategies in the very large mass range displayed in Fig 2, supports the idea that selection mechanisms should operate in terms of equal fit according to Darwinian evolution.

In the fundamental theorem of natural selection, Fisher (1930) states that the rate of increase in mean fitness (caused by natural selection; Price 1972) is equal to the genetic variance of a species, conceptually linking natural selection with Mendelian genetics. Natural selection can only increase fitness by reducing genetic variance (i.e., selecting away undesirable alleles; Basener & Sanford 2018), and thus, without mutations and given enough time, selection must reduce genetic variance all the way to zero and fitness must reach a maximum, according to Fisher’s theorem and as confirmed through simulations (Basener & Sanford 2018). The ‘Equal Fitness Paradigm’ seems to suggest that a maximum fitness value has been reached by the coexisting species in the current conditions.

Additionally, Demetrius (1974, 1975) found a relation between ‘evolutionary’ entropy (of a population) and reproductive potential with fitness (measured by the Malthusian parameter), in which Fisher’s theorem is also obtained as a corollary for a Hardy-Weinberg equilibrium. In Demetrius’s formalism, the Malthusian parameter is analogous to the Gibbs free energy (Demetrius 1997) and also, behave effectively like a thermodynamic potential: being the maximum (potential) growth that can be performed by a given population, that becomes null when such population reaches abundance equilibrium. In thermodynamic theory, Gibbs free energy is minimized when entropy is maximized and according to the second law of thermodynamics, maximum entropy is found only in the state of thermodynamic equilibrium, where (the theorem of) equipartition of energy holds. The ‘Energetic Equivalence Rule’ (Damuth 1981), which was recently verified for ~3000 species in Hatton et al. (2019), seems to be a version of such an equipartition of energy in living matter and thus is also consistent (under abundance equilibrium) with being in the state in which maximum entropy is reached.

Another relation for r_m_ that can be derived using the definition of G (Eq. 6) is the relation that predicts an inverse correlation with the generation time t_gen_:

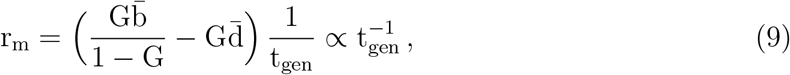

and this is also a relation that can be tested. For this purpose, we use the data in Duncan et al. (1997), which compiles the ages at first reproduction (~ t_gen_) and maximum population growth rates for mammals. Fig 3 displays the data collected in Duncan et al. (1997), which is in overall agreement with Eq. 9, with no relevant slope with body mass. The solid line denotes the best fit to the data (r_m_t_gen_ = 1.38 M^−0.067^), which has an average scatter of 0.2 dex. Dillingham et. al (2016) also found equivalent results on this expected inverse relationship using a more sophisticated Bayesian analysis.

**Fig. 3.**
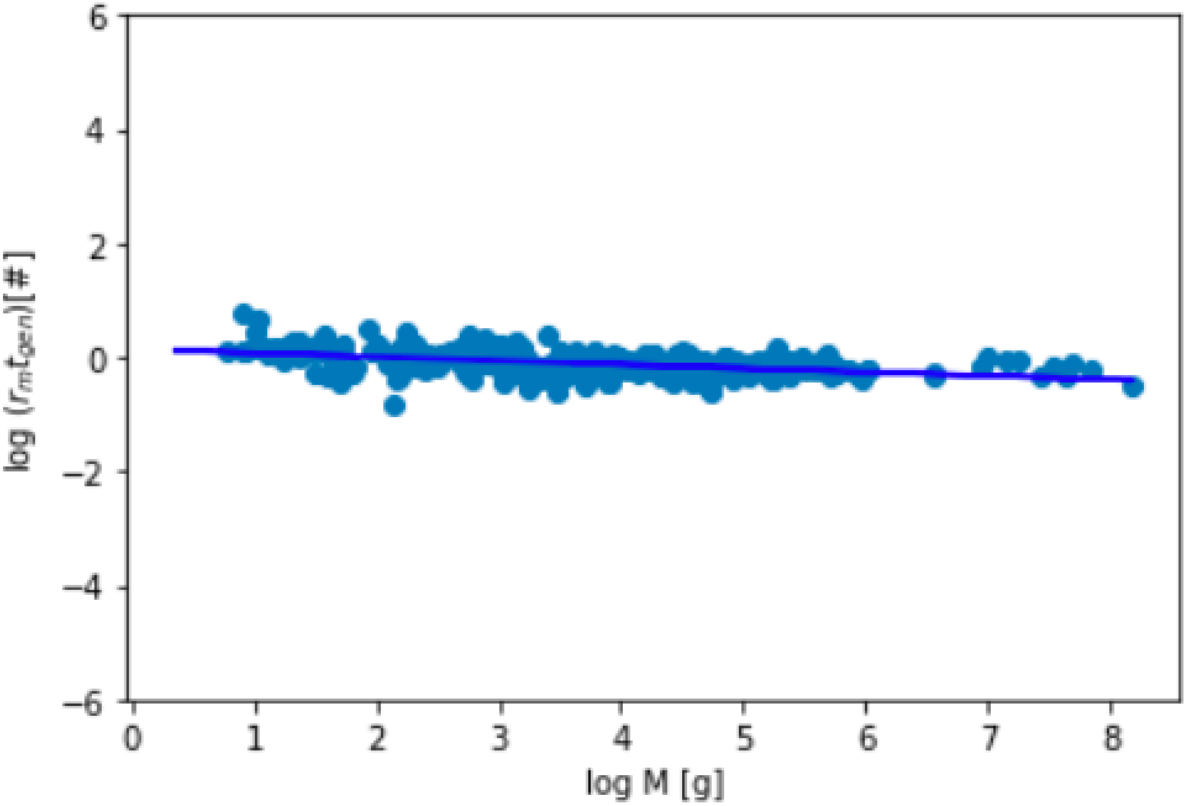
Generation times (t_gen_) times the maximum population growth rates (r_m_) for mammals, compiled by Duncan et al. (1997). The solid line denotes r_m_t_gen_ = 1.38 M^−0.067^, which is the best fit to the data and has an average scatter of 0.2 dex. The best-fitted relation is in overall agreement with the predicted Eq. 9, with no relevant slope with body mass.

The normalizations from the best fit to the data in Figs 2 & 3 give estimations for r_m_t_life_ ~ 3.35 (Fig 2) and r_m_t_gen_ ~ 1.38 (Fig 3), which in principle can be combined with Eq. 8 & 9 to determine some of their constants. Unfortunately, it is not possible to determine the constants 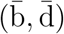 associated with the dimensionless birth and death rates since Eq. 8 & 9 are not (linearly) independent equations, but it is still possible to determine that 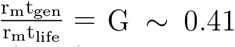. This is for the average lifespan in the wild because in Hatton et al (2019), lifespans were normalized to those values, and since maximum lifespans are about 2.5 times the average lifespans in the wild (McCoy & Gillooly 2008), we determine from Figs 2 & 3 that 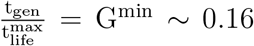, which agrees with the value determined from the average of the predicted G values in Table 2 (G^min^ ~ 0.17).

## 5. Summary

In this paper, we explored the implications of the new metabolic relation (Escala 2019) for ontogenetic growth and found that it can be described by a universal ontogenetic growth curve for all studied species without the aid of ad hoc fitting parameters. The same universal growth curve in West et al (2001) and Banavar et al (2002) is found when certain dimensionless quantities are properly defined, but in our case, the characteristic growth timescale (t_growth_) is set by the inverse of the heart frequency 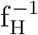, rescaled by the ratio between the specific energies (per unit of mass) required for metabolism and the creation of new biomass; these are all quantities with clear physical or biological meaning and not obscure fitting parameters. The results for universal growth have the same interpretations as before in terms of conservation of energy, since they come from the same Bertalanffy-type equation, but in our case, the results do not rely on a specific model. Instead, they illustrate the advantages obtained when empirical data are properly described and quantified, reaching this simple description without extra assumptions.

We also explored the implications of the discussed ontogenetic growth model for the generation time, finding that the predicted t_gen_ can explain the origin of several ‘Life History Invariants’ when is it combined with the invariant number of respiration cycles per lifetime, a relation that comes from the generalization of the well-known relation of the constant number of heartbeats in a mammal’s lifetime (Levine 1997; Cook et al. 2006). In particular, regarding the invariant ratio between the lifespan and age at maturity 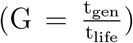, which has been traditionally explained in terms of life-history evolution theory, in our formalism, the value of this life-history invariant is predicted in terms of the relevant energetics and the invariant number of respiration cycles per lifetime. The predicted G and its variation between taxonomic groups shows consistency with the empirically determined value (Table 2). We also showed that other life-history invariants (Charnov 1993) come directly from the universal ontogenetic growth curve and the invariant G.

We finally studied predictions for population growth, finding that the invariant G implies a Malthusian parameter r_m_, or intrinsic population growth rate, that is inversely proportional to both t_gen_ and t_life_. We find that these inverse relations are indeed observed in nature, with no relevant slope with body mass and relatively low scatter (0.5 dex for r_m_t_life_ in Fig 2 and 0.2 dex for r_m_t_gen_ in Fig 3). The ratio 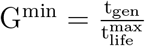 that can be estimated from the two best-fitted relations 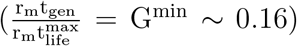 is also consistent with our predicted G^min^ (~ 0.17; Table 2). We find relatively constant values for the total births and deaths per capita per characteristic time (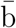 and 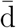 in Eq. 8), besides the diversity of life histories in living organisms on Earth, which supports the so-called ‘Equal Fitness Paradigm’.

In our formalism, the allometric scaling relations for several critical timescales and rates (t_life_, t_gen_ & r_m_) are all derived from their proportionality to f_H_; therefore, the mass scaling f_H_ ∝ M^−1/4^ directly explains other well-known allometries (lifespan, generation-time and Fenchel allometries), naturally explaining why most biological times and rates scale as M^−1/4^ and M^1/4^ (Savage et al. 2004, Burger et al. 2021). Additionally, the variations from 1/4 scaling should be explained in terms of the variation in the f_H_ mass scaling, since such quantities (t_life_, t_gen_ and r_m_) should have the same mass scaling as f_H_ regardless of whether they are 1/4, as was the case for the metabolic rate relation (Escala 2019).

Nevertheless, our formalism is empirically motivated and does not explain why f_H_ should scale as M^−1/4^ or with another exponent; thus, it is compatible with previous attempts that explained the exponent for the allometric scaling of metabolism and related quantities (West et al 1997, 1999; Banavar et al 1999, 2010; Darveau et al 2002) insofar as the mechanism is based on the anatomy and physiology of the circulatory system. The difference between these approaches is similar to the difference in physics between the (empirically-based and axiomatic) laws of thermodynamics and the statistical mechanics that explain them; in this case, the new metabolic rate and the invariant number of respiration cycles per lifetime are analogous to the thermodynamic laws, and such a synthesis is also required for the formulation of a general theory of biodiversity (Marquet 2017).

Finally, it is important to emphasize that our predictions were successfully tested without the aid of any free (fitting) parameters. The fitting procedures are shown in Figs 1, 2 & 3 only for comparison with our predictions, not to determine any free parameters for our formalism. Thus, our formalism fulfill the criteria suggested by Ginzburg & Jensen (2004) for judging ecological theories, in terms of reducing to a minimum the number of parameters with no empirically determined range. Moreover, the constants that define the critical relations for metabolism (E_2019_) and lifespan (N_r_) are successfully tested further in Fig 1 (E_2019_) and in terms of the G value determined in Figs 2 & 3 (E_2019_ and N_r_; Eq. 6), showing that the two simple relations for metabolism and lifespan used in this paper can successfully explain a variety of complex phenomena. This resembles the reductionism seen in the physical sciences, where simple laws account for multitudes of complex phenomena.

